# *Pkd1* mutation has no apparent effects on peroxisome structure or lipid metabolism

**DOI:** 10.1101/2021.02.08.430145

**Authors:** Takeshi Terabayashi, Luis F Menezes, Fang Zhou, Hongyi Cai, Peter J Walter, Hugo M Garraffo, Gregory G Germino

**Affiliations:** Kidney Disease Branch; National Institutes of Diabetes and Digestive and Kidney Disease, NIDDK, National Institutes of Health, Bethesda, MD, United States; Clinical Mass Spectrometry Core; National Institutes of Diabetes and Digestive and Kidney Disease, National Institutes of Health, Bethesda, MD, United States

## Abstract

**Background:** Multiple studies of tissue and cell samples from patients and pre-clinical models of autosomal dominant polycystic kidney disease report abnormal mitochondrial function and morphology and suggest metabolic reprogramming is an intrinsic feature of this disease. Peroxisomes interact with mitochondria physically and functionally, and congenital peroxisome biogenesis disorders can cause various phenotypes, including mitochondrial defects, metabolic abnormalities and renal cysts. We hypothesized that a peroxisomal defect might contribute to the metabolic and mitochondrial impairments observed in autosomal dominant polycystic kidney disease.

**Methods:** Using control and *Pkd1*^*-/-*^ kidney epithelial cells, we investigated peroxisome abundance, biogenesis and morphology by immunoblotting, immunofluorescent and live cell imaging of peroxisome-related proteins and assayed peroxisomal specific β-oxidation. We further analyzed fatty acid composition by mass spectrometry in kidneys of *Pkd1*^*fl/fl*^; *Ksp-Cre* mice. We also evaluated peroxisome lipid metabolism in published metabolomics datasets of *Pkd1* mutant cells and kidneys. Lastly, we investigated if the C-terminus or full-length polycystin-1 co-localize with peroxisome markers by imaging studies.

**Results:** Peroxisome abundance, morphology and peroxisome-related protein expression in *Pkd1*^*-/-*^ cells were normal, suggesting preserved peroxisome biogenesis. Peroxisomal β-oxidation was not impaired in *Pkd1*^*-/-*^ cells, and there was no obvious accumulation of very long chain fatty acids in kidneys of mutant mice. Re-analysis of published datasets provide little evidence of peroxisomal abnormalities in independent sets of *Pkd1* mutant cells and cystic kidneys, while providing further evidence of mitochondrial fatty acid oxidation defects. Imaging studies with either full length polycystin-1 or its C-terminus, a fragment previously shown to go to the mitochondria, showed minimal co-localization with peroxisome markers.

**Conclusions:** Our studies showed that loss of *Pkd1* does not disrupt peroxisome biogenesis nor peroxisome-dependent fatty acid metabolism.

**Key points:**

- While mitochondrial abnormalities and fatty acid oxidation impairment have been reported in ADPKD, no studies have investigated if peroxisome dysfunction contributes to these defects.
- We investigated peroxisome morphology, biogenesis and function in cell and animal models of ADPKD and investigated whether polycystin-1 co-localized with peroxisome proteins.
- Our studies show that loss of *Pkd1* does not disrupt peroxisome biogenesis nor peroxisome-dependent fatty acid metabolism.

## Introduction

Autosomal dominant polycystic kidney disease (ADPKD) is one of the most common genetic diseases, affecting ∼1/1000. It is primarily caused by mutations in the *PKD1* or *PKD2* genes and is characterized by the appearance and growth of cystic lesions in multiple organs, particularly kidney and liver. ADPKD often results in end stage renal failure (ESRF) by the sixth decade, as the currently available therapy provides only limited benefit.

A growing body of evidence suggests metabolic reprograming is an intrinsic feature of ADPKD [1-3]. Multiple groups have investigated and reported mitochondrial functional or morphological abnormalities in experimental ADPKD models and human patient samples, including impaired glucose metabolism [2], fatty acid oxidation [4], and disorganized mitochondrial cristae with altered mitochondrial network [5, 6]. While a functional link between metabolism and the gene products of *PKD1* (polycystin-1; PC1) and *PKD2* (polycystin-2; PC2) is not yet clear, there have been reports of mitochondrial targeting of a cleavage product of PC1 [6], and PC2 has been proposed to regulate mitochondrial calcium homeostasis [7, 8].

A less well studied potential cellular mechanism for metabolic dysregulation in ADPKD is peroxisomal dysfunction. Peroxisomes are single-membrane lined organelles present in virtually all eukaryotic cells and crucial for several metabolic pathways, including synthesis of plasmalogen and other glycerophospholipids, long fatty acid β-oxidation, and metabolism of cholesterol and bile acids [9]. Mitochondria and peroxisomes interact both physically and functionally [10] [11]. In fact, while most peroxisomes are produced through growth/division cycles, *de novo* peroxisome biogenesis requires fusion of pre-peroxisome vesicles derived from both the ER and mitochondria [12]. The mitochondria/peroxisome functional overlap includes fatty acid oxidation, response to oxidative stress and shared components in organelle fission machinery [11].

Genetic peroxisomal disorders often manifest through progressive metabolic dysfunction and developmental defects, such as craniofacial and ocular malformations, severe central neurological phenotypes, and liver and kidney abnormalities, including small renal cysts [9, 13, 14]. The mechanisms leading to these abnormalities is still poorly characterized, but it is noteworthy that an orthologous mouse model of Zellweger syndrome (a peroxisome biogenesis disorder) had multiple mitochondrial abnormalities including large aggregates of pleomorphic mitochondria with abnormal cristae [15] and altered dynamics leading to increased fragmentation [16]. A partially penetrant mitochondrial phenotype was also reported in human patients [17], leading to the suggestion that mitochondrial dysfunction secondary to peroxisome defects contributes to the pathogenesis of Zellweger syndrome [16].

Given this physiological and pathophysiological overlap between peroxisomes and mitochondria (Table 1), we set out to investigate whether PC1 is primarily involved in peroxisomal activity and if a peroxisomal defect might contribute to the mitochondrial defects observed in ADPKD.

**Table 1.**
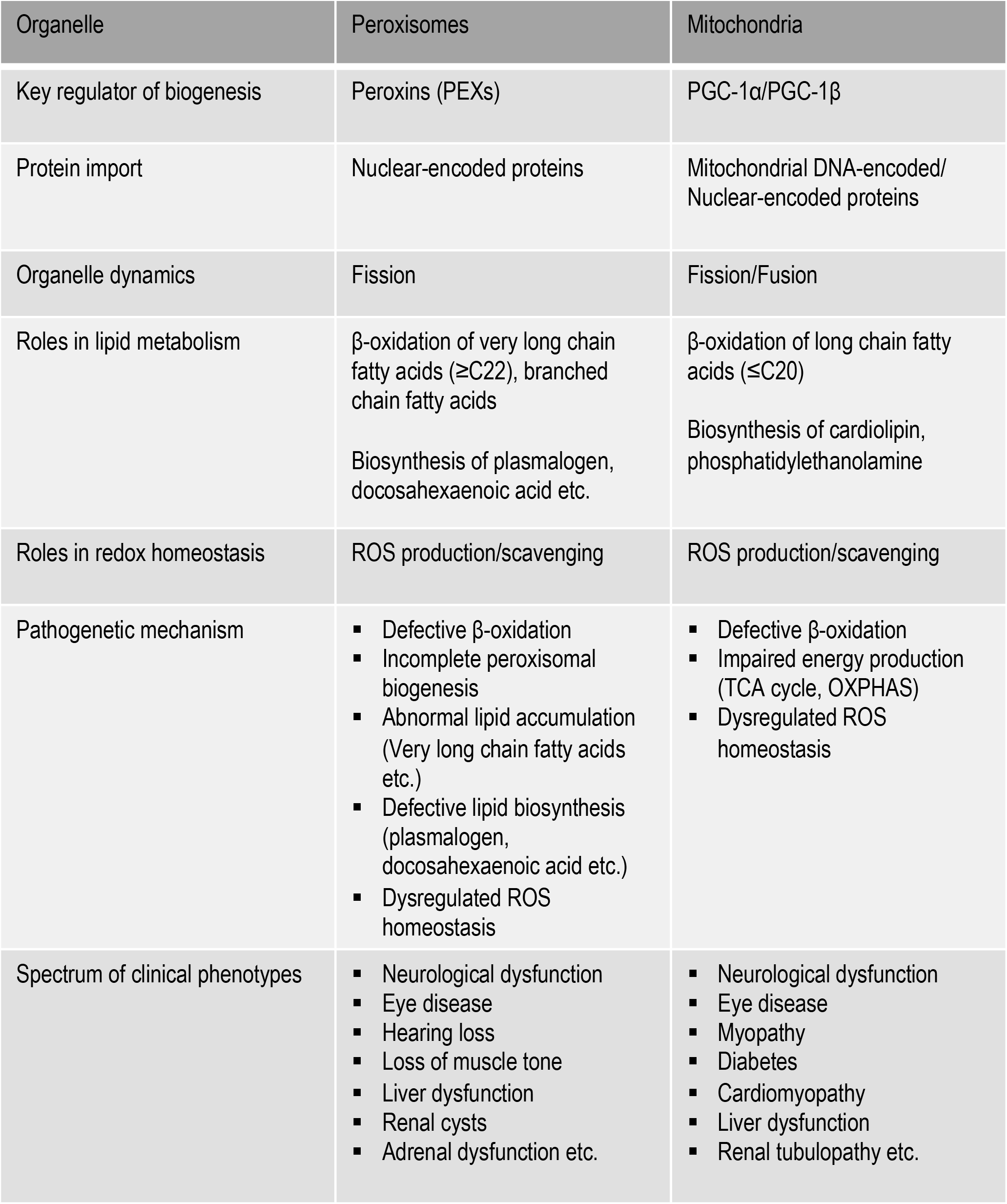
Physiological and pathophysiological interaction between peroxisomes and mitochondria

## Results

### Evaluating peroxisome biogenesis pathways in Pkd1 mutant kidney epithelial cells

Peroxisomal disorders can be broadly divided into two groups: those that are the result of biogenesis defects, a genetically and phenotypically heterogeneous group mostly characterized by peroxisomal protein import deficiency; and those resulting from single enzyme defects and mostly impacting specific metabolic pathways [14]. A common feature of both is peroxisomes larger in size but reduced in numbers (down to about 20% of normal) [18]. The *de novo* biogenesis of peroxisomes is a stepwise process starting with the assembly of the import machinery, primarily Pex3 and Pex16, which subsequently import other membrane proteins bound to Pex19 [19] [20] [21]. Careful characterization of this process suggests that Pex3 localized in mitochondria (stage 0) accumulate in pre-peroxisomal vesicles (stage Ia) which merge with ER-derived vesicles containing Pex16 (stage Ib), next importing additional membrane proteins, such as PMP70 (stage II) and finally matrix proteins, such as catalase (stage III), forming mature peroxisomes [12] (Figure 1a).

**Figure 1.**
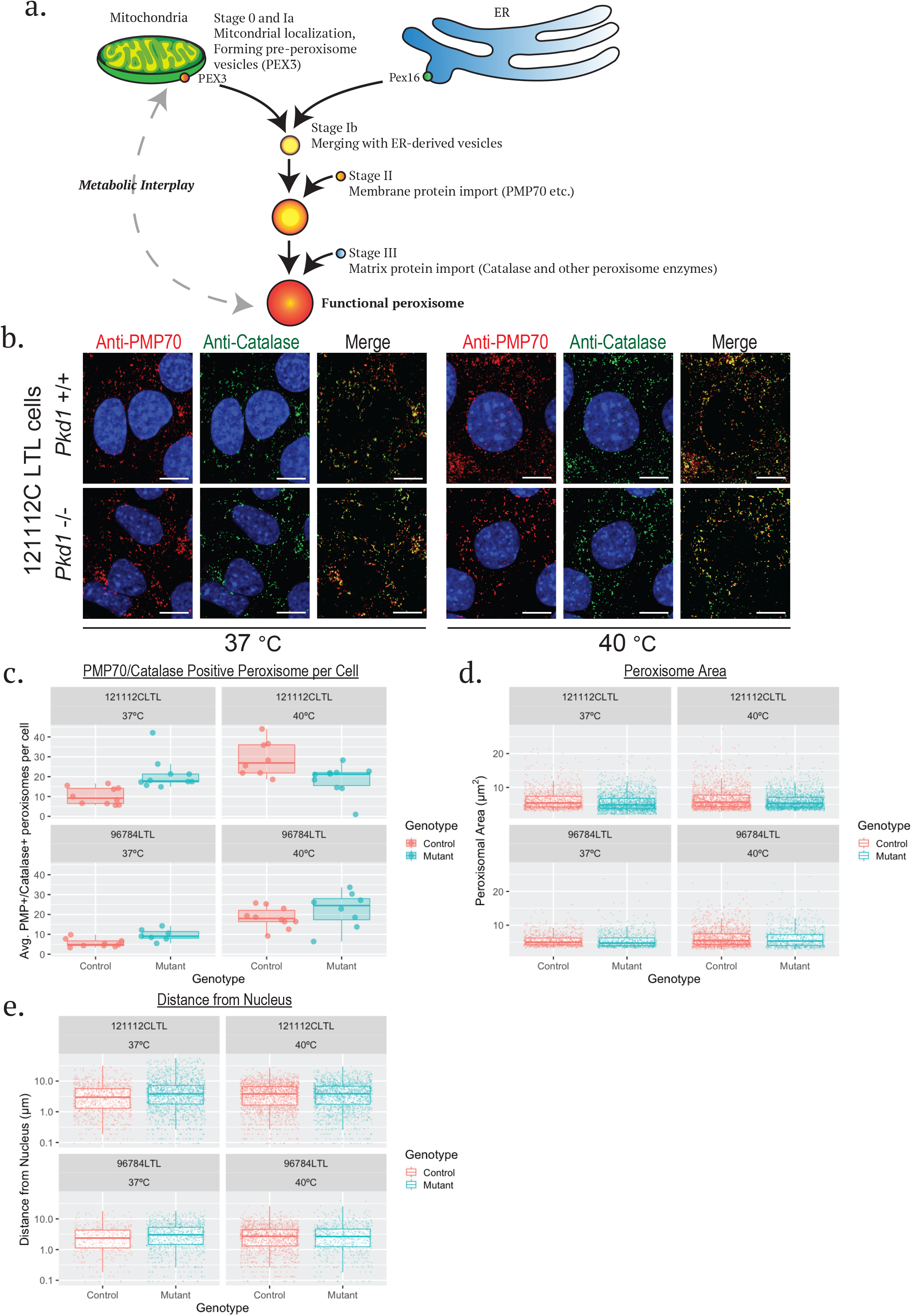
Peroxisomal biogenesis is preserved in *Pkd1* mutant kidney epithelial cells. a) A schematic diagram of the current model for *de novo* formation of peroxisomes showing the hybrid origin of peroxisomes: nascent peroxisome vesicles derived from mitochondria (PEX3) and ER (PEX16 and other peroxins) fuse and then undergo subsequent stepwise assembly of peroxisome components to form mature peroxisomes. The functional peroxisomes interplay with mitochondria in metabolic cellular processes such as fatty acid oxidation. b) Representative images of *Pkd1* control and mutant kidney epithelial cells (121112CLTL cell line) double immunostained for PMP70 and catalase. Peroxisomal number, size and distribution were similar in *Pkd1* mutant and control cells both at 37 °C (the images in the top two rows) and after 48 hours incubation at 40 °C (the images in the lower two rows). Scale bar, 10µm. c) Boxplots showing quantification of the number of PMP70/catalase double positive peroxisomes in *Pkd1* mutant and control cells derived from two kidney epithelial cell lines (121112CLTL, 96784LTL) cultured at 37°C and at 40 °C. None of the cell lines manifest a significant decrease in peroxisomal number at high temperature (40°C). d, e) Boxplots show quantification of peroxisomal area (d) and distance between PMP70 positive peroxisomes to nucleus (e) in the 121112CLTL and 96784LTL kidney cell lines.

In peroxisomal biogenesis diseases, numerous PMP70 positive vesicles (stage II) fail to import matrix proteins, resulting in reduced numbers of mature/functional peroxisomes [22]. Therefore, peroxisome number, shape, localization and protein composition are closely related to their function [14, 23]. To investigate whether peroxisome biogenesis pathways are affected by loss of PC1 function, we counted the number and size of PMP70/catalase positive vesicles in two pairs of *Pkd1* mutant and control kidney epithelial cell lines. We found that the size, number and distribution of vesicles was similar in both genotypes (Figure 1b, panels in left three columns; Figure 1c, left panels). It has been reported that a cellular phenotype cannot be detected in patient-derived fibroblasts of individuals with milder forms of peroxisomal diseases under normal conditions, but abnormal catalase import into peroxisomes can be observed in cells cultured at high temperatures (40°C) [24, 25]. To address this possibility, we investigated catalase staining patterns in *Pkd1* mutant and control kidney epithelial cells stressed with high temperature. We found a similar punctate pattern in both genotypes (Figure 1b, panels in right three columns), and in quantifying the results we observed no decrease in the number and size or significant change in the location of mature peroxisomes (i.e. PMP70; catalase double positive vesicles) in any of the cell lines after incubation for 48 hours at 40°C (Figure 1c-e), consistent with preserved peroxisome biogenesis.

A variety of models have been proposed for peroxisome biogenesis, but they mostly involve two main processes: protein cycling between cytoplasm and peroxisomes, and changes in peroxisomal protein stability [26]. We checked protein abundance by western blotting as a crude readout for stability and detected no genotype-specific pattern in Pex3, Pex5 and PMP70 expression in *Pkd1* mutant cells (Figure 2 and Supplementary Figure S1).

**Figure 2.**
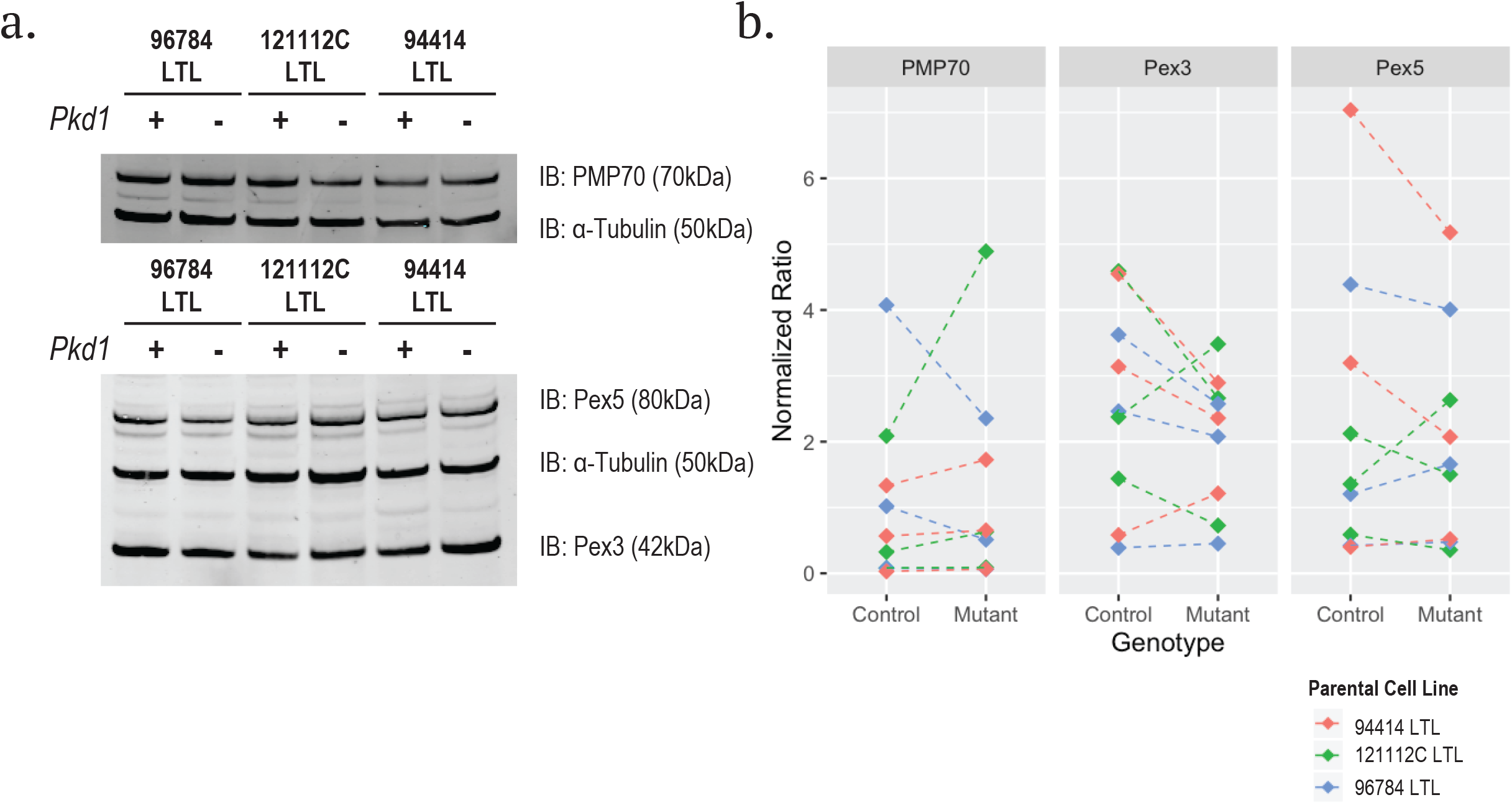
Expression of peroxisomal associated proteins in *Pkd1* control and mutant kidney epithelial cells. a, b) Representative immunoblot of protein lysates from three different wild type (*Pkd1+*) and mutant (*Pkd1-*) pairs of kidney cell lines probed for peroxisomal associated proteins PMP70 (a), Pex3 and Pex5 (b). α-tubulin was used as a loading control. b) Quantification of the data from the immunoblot studies shows comparable levels of protein expression in *Pkd1* control and mutant cells. The signal intensity values of PMP70, Pex3 and Pex5 were individually normalized to the intensity of the α-tubulin band detected in the same lane. The results for three separate studies of each cell line pair are shown.

Peroxisome homeostasis also involves its dynamic morphological regulation of elongation, constriction and division (fission) [27], therefore morphological change such as elongated/enlarged peroxisomes has been implicated in the pathogenesis of peroxisomal disorders [25]. For assessment of peroxisomal morphology, we performed live cell imaging using a peroxisome-import reporter to target EGFP tethered with a C-terminal peroxisomal targeting sequence [EGFP-SKL; [28]] in transfected cells. We found that both *Pkd1* wild type and mutant cells exhibited punctate patterns of EGFP signal and that peroxisomal area did not differ based on genotype (Figure 3a, b). These data are consistent with the catalase/PMP70 results in Figure 1 and suggest protein import into peroxisome matrix is likely intact in *Pkd1* mutant kidney epithelial cells. We also did not detect genotype-dependent differences in shape (as measured by ellipticity) [29], suggesting peroxisomal morphology was not affected by *Pkd1* genotype (Figure 3c).

**Figure 3.**
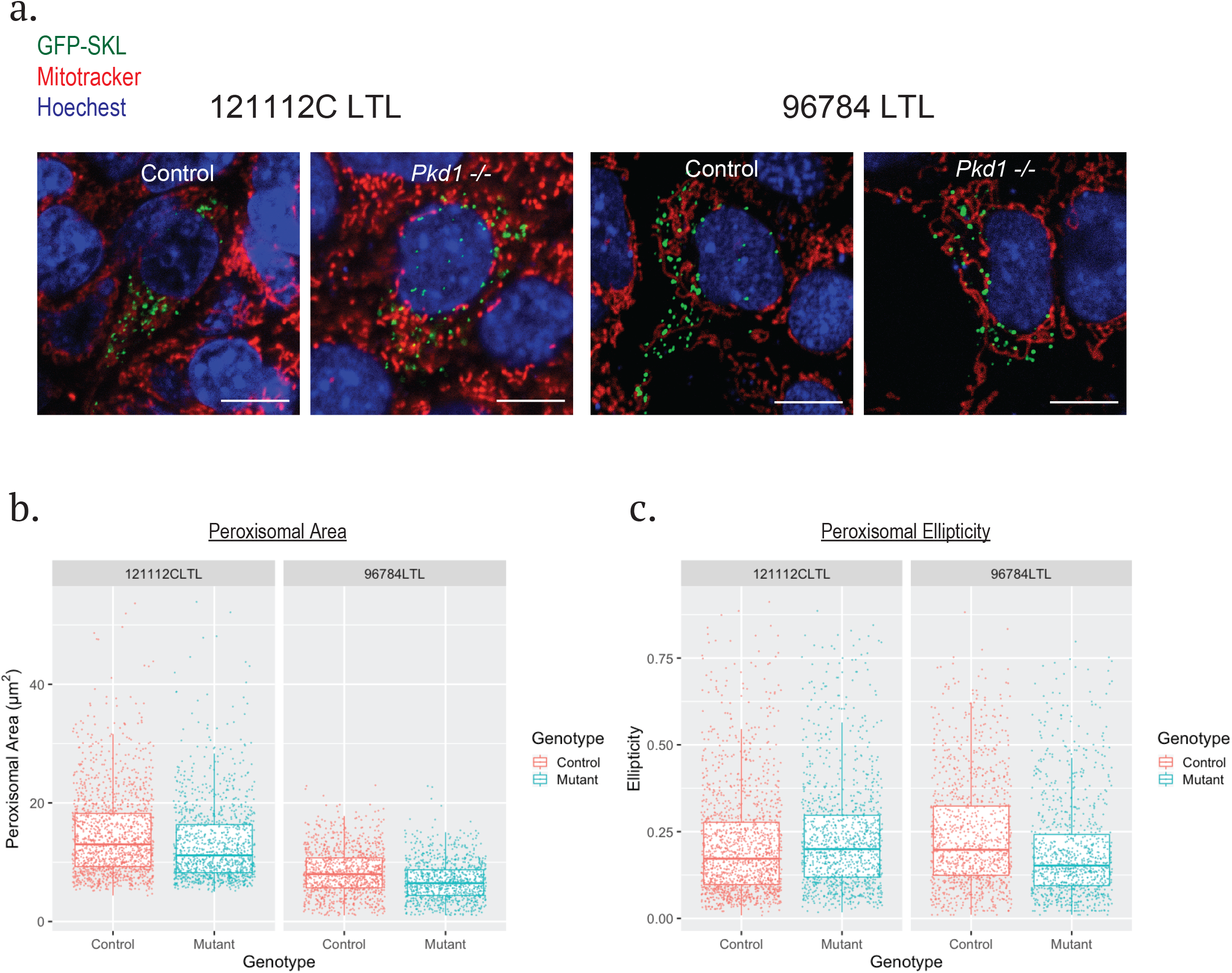
Assessment of peroxisomal matrix protein import machinery and morphology by live cell imaging in *Pkd1* control and mutant kidney epithelial cells. a) Representative images of live cell imaging of kidney epithelial cells expressing EGFP-SKL. The *Pkd1* control and mutant cells show a robust, similar punctate pattern of fluorescent signals. Scale bar, 10 µm, b, c) Boxplots show quantification of the area (b) and peroxisomal ellipticity (c) for approximately 1000-1400 SKL-EGFP positive peroxisomes/cell line-genotype for two pairs of *Pkd1* control and mutant kidney epithelial cell lines. The results show that *Pkd1* mutant peroxisomes have normal matrix import activity and morphology.

### Investigating peroxisomal β-oxidation in Pkd1 mutant cells and tissues

While morphological abnormalities in peroxisomes is a common feature of peroxisomal diseases, in some milder forms or in diseases with defects in specific enzymatic pathways, peroxisomes can appear normal [14]. Peroxisomes especially play a key role in fatty acid oxidation and are the organelles exclusively responsible for oxidation of very long chain fatty acids (VLCFA) such as hexacosanoic acid (C26:0) and tetracosanoic acid (C24:0) or the beta-oxidation of C24:6 to C22:6 (Figure 4a) [30] [31]. A decrease in VLCFA oxidation also results in an increase in substrate for further fatty acid chain elongation [32]. Therefore, screening for decreased oxidation rate of VLCFA or accumulation of VLCFA is a reliable method for identifying peroxisomal dysfunction [31] (Figure 4b).

**Figure 4.**
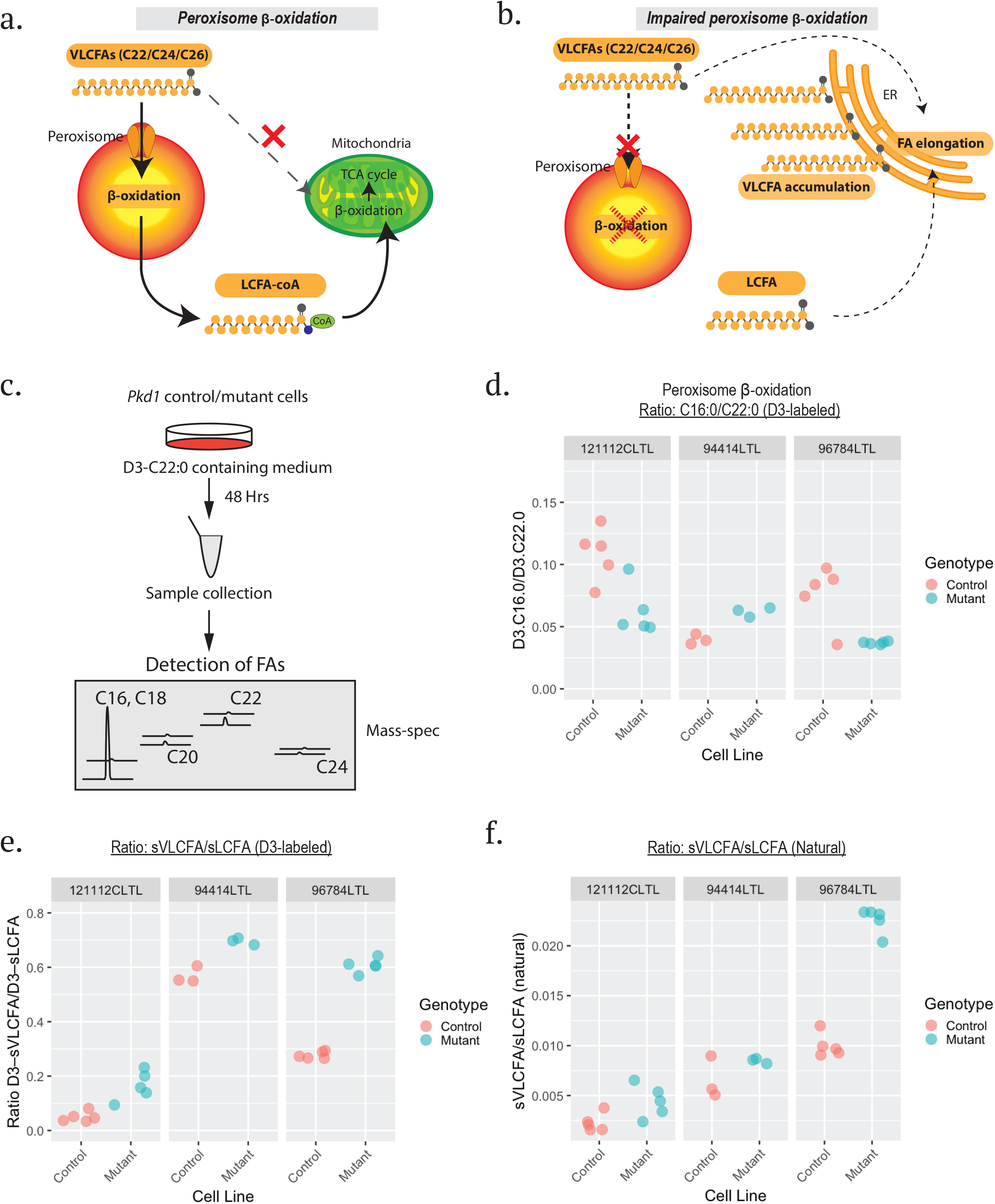
Peroxisome β-oxidation in *Pkd1* control and mutant kidney epithelial cells. a, b) Schematic diagrams illustrating the unique role of the peroxisome in fatty acid metabolism and consequences of its disruption. VLCFA cannot enter the mitochondria and must be converted to LCFA by β-oxidation in the peroxisome prior to further metabolism in mitochondria (a). When peroxisomal function is compromised, the rate of VLCFA oxidation is decreased, increasing the pool of LCFA substrates available for fatty acid elongation leading to further accumulation of VLCFA (b). c) A diagram showing a workflow for measurement of peroxisome β-oxidation using deuterium-labeled docosanoic acid (D3–C22:0). “FA” indicates “fatty acids”. d) A scatter plot presenting the ratio of deuterium-labeled (D3–) C16:0/D3–C22:0 (c) as readout for peroxisome β-oxidation in three distinct pairs of control/*Pkd1* mutant mouse kidney epithelial cell lines. The results indicate no genotype-dependent alteration in the peroxisomal oxidation rates. e, f) Scatter plots showing the ratio of saturated VLCFA (sVLCFA) to saturated long chain fatty acid (sLCFA) of D3-labeled species: (D3–C24:0 and D3–C26:0)/(D3–C16:0, D3–C18:0 and D3– C20:0) (e), and natural fatty acids: (C24:0 and C26:0)/(C16:0, C18:0 and C20:0). The raw datasets are available in Supplementary table 1.

We used a number of strategies to assess peroxisomal function in *Pkd1* mutant cells and tissues by mass spectrometry. We first quantified long chain fatty acid (LCFA) levels and VLCFA levels in multiple replicates (N=3-5) of three pairs of matched control and *Pkd1* mutant cell lines treated with stable isotope labeled VLCFA (deuterium-labeled docosanoic acid: D3– C22:0) (Figure 4c) [33]. By measuring both natural and labeled fatty acids, we assessed both steady-state and dynamic patterns of oxidation or elongation (Supplementary Table 1). We found that the ratio of D3–C16:0/C22:0 did not show significant genotype-dependent difference, therefore indicating comparable peroxisome β-oxidation rates in *Pkd1* mutant cells (Figure 4d). The results for the VLCFA/LCFA analyses were less straight-forward. The ratio of labeled VLCFA/LCFA was increased in each of the three *Pkd1* mutant cell lines compared to its paired control, suggesting a possible increase in the rate of fatty acid chain elongation (Figure 4e). In contrast, there was no genotype-dependent difference in the ratio of unlabeled VLCFA/LCFA in two out of the three cell lines, indicating that in the steady state there was not a consistent pattern of net accumulation of VLCFAs (Figure 4f).

To further examine this question, we compared fatty acid levels in cystic and normal mouse kidneys. We used the *Pkd1*^*fl/fl*^; *Ksp-Cre* mouse model in which *Pkd1* is conditionally inactivated in epithelial cells by cadherin-16-driven cre expression [34], and we grouped the data for the *Pkd1*^*fl/+*^*;Ksp-Cre* positive and *Ksp-Cre* negative samples for analyses since the kidneys are phenotypically normal in mice of either genotype (Figure 5). We selected post-natal days 3 and 6 since these timepoints demark an interval when the kidneys are transitioning from mildly to moderately cystic and measured C10-C24 fatty acids (Figure 5).

**Figure 5.**
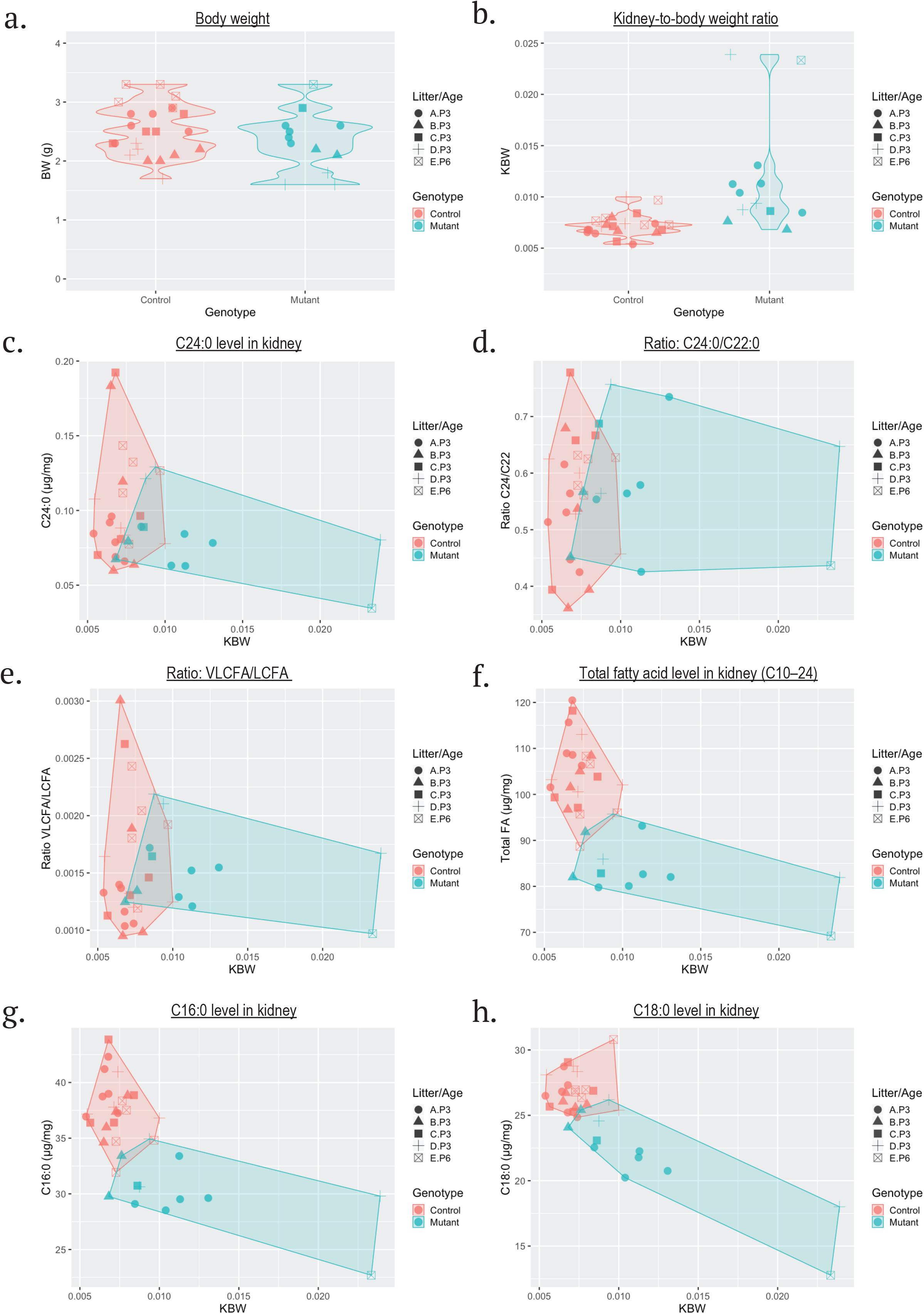
Fatty acid profile in kidneys of *Pkd1 control and mutant mice*. a, b) Violin plots show comparisons of body weight (a) and kidney to body weight ratio (KBW) (b) between control (*Pkd1*^*fl/+*^*;Ksp-Cre* positive and *Ksp-Cre* negative) and mutant mice (*Pkd*^*fl/fl*^*;Ksp-Cre*). c) A scatter plot shows the kidney-body weight ratio (KBW) (x-axis) and C24:0 level in mouse kidneys (y-axis). d) A scatter plot shows KBW (x-axis) and ratio of C24:0/C22:0 (y-axis). e) A scatter plot presenting KBW (x-axis) and the ratio of VLCFA (C24:0 and C26:0) to LCFA (C16:0, C18:0 and C20:0) in mice kidneys (y-axis) suggests no significant differences between *Pkd1* control and mutant samples. f) A scatter plot presenting correlation between KBW (x-axis) and level of total fatty acids (C10– C24) (y-axis) reveals distinct clusters of *Pkd1* control and mutant mice samples suggesting genotype-based differences. g, h) Scatter plots showing correlation between C16:0 (g, y-axis) or C18:0 levels (h, y-axis) and KBW (x-axis) reveal genotype-dependent clusters. The raw datasets are available in Supplementary Table 2.

As expected, there was no difference in body weight between the groups, while kidney/body weight ratios, an indirect measure of cystic enlargement, were generally greater in the mutants with higher ratios in older mutants (Figure 5a, b). We found that the levels of C24:0 trended lower in cystic kidneys (*Pkd1*^*fl/+*^*;Ksp-Cre* and *Ksp-Cre* negative 0.10±0.04 μg/mg vs *Pkd1*^*fl/fl*^*;Ksp-Cre* 0.08±0.03 μg/mg, p=0.12; Figure 5c) while the C24:0/C22:0 ratio [13] was no different (*Pkd1*^*fl/+*^*;Ksp-Cre* and *Ksp-Cre* negative 0.56±0.11 vs *Pkd1*^*fl/fl*^*;Ksp-Cre* 0.58±0.11, p=0.53, Figure 5c), indicating that there was a broad decrease in levels of VLCFA. Likewise, we found no difference between normal and cystic kidneys in the ratio of VLCFA to LCFA (*Pkd1*^*fl/+*^*;Ksp-Cre* and *Ksp-Cre* negative 0.0015±0.0006 vs *Pkd1*^*fl/fl*^*;Ksp-Cre* 0.0015±0.0004, p=0.98, Figure 5d), indicating no obvious accumulation of VLCFA in the orthologous model of ADPKD. We found, however, that the tissue concentrations of total fatty acids (C10–C24 normalized to kidney weight) were lower in cystic kidneys compared to controls (*Pkd1*^*fl/+*^*;Ksp-Cre* and *Ksp-Cre* negative 104.6±7.7 μg/mg vs *Pkd1*^*fl/fl*^*;Ksp-Cre* 83.9±7.1 μg/mg, p<6.3e-9; Figure 5f). While this result is at least explained in part by the dilutional effects of cystic fluid contributing to total kidney weight (Supplementary Table 2), we can’t exclude a metabolic cause since some LCFA were also lower in *Pkd1* mutant MEFs (Figure 6e-g). We also confirmed that the differences could not be explained by sex effects as there were approximately equal numbers of male and female samples in each set and there was no difference between male and female values within a genotype (Supplementary Figure S2a).

**Figure 6.**
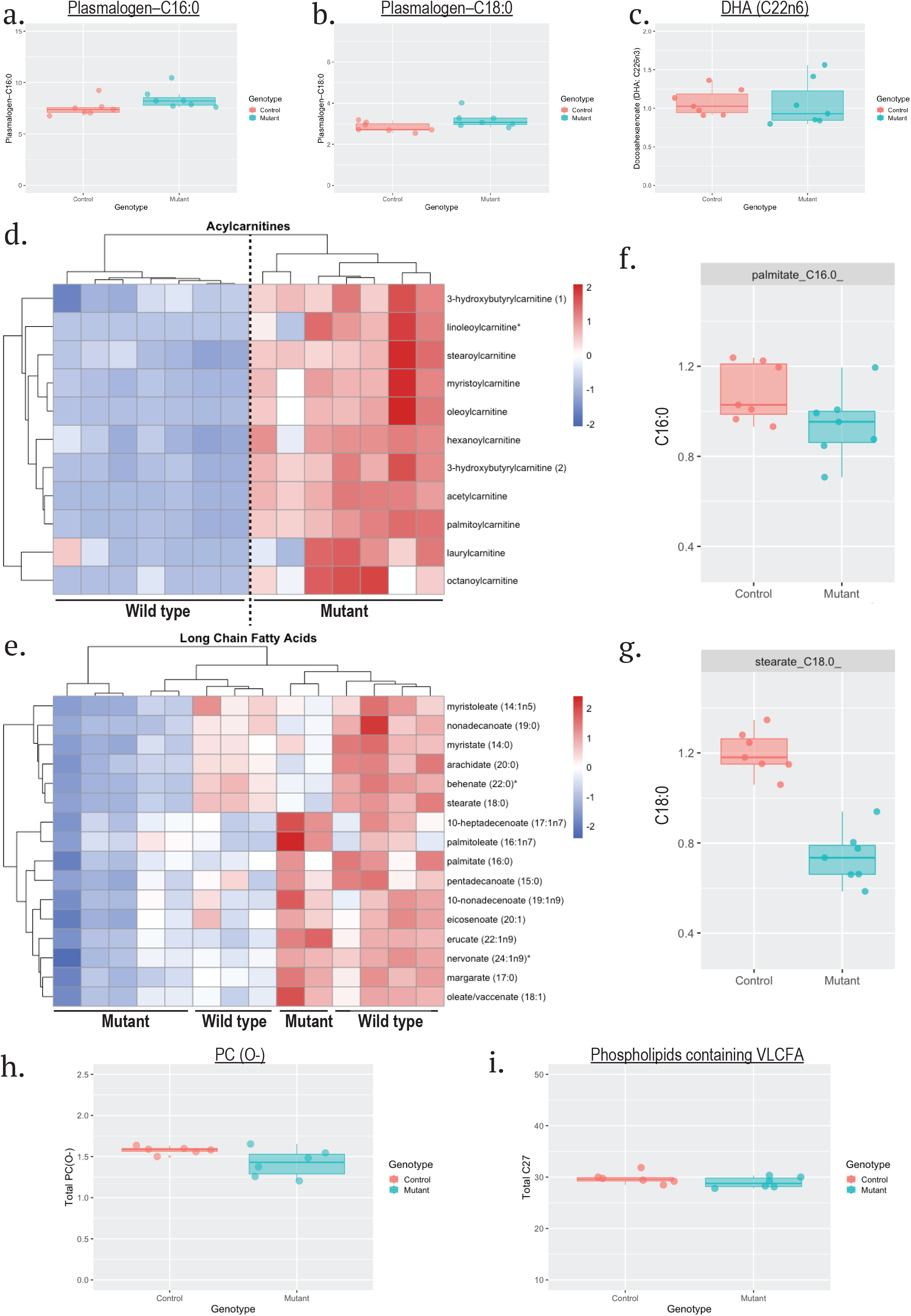
Re-analysis of published metabolomic and lipidomic datasets of *Pkd1* mutants. a–g) Analysis of published dataset of lipid metabolites detected in control and *Pkd1* mutant mouse embryo fibroblasts (MEF) [35]. a–c) Boxplots showing levels of peroxisome-associated lipids: plasmalogen–C16:0 (a), plasmalogen–C18:0 (b), and docosahexaenoic acid (c) respectively. Values for each compound were normalized by protein concentration (Bradford assay) followed by rescaling to set the median equal to 1 [35], then values were totaled for plasmalogen–C16:0 and plasmalogen– C18:0. d) A heatmap showing relative abundance of acylcarnitines in control and *Pkd1* mutant MEFs. Clustering shows clear separation based on *Pkd1* genotype suggesting acylcartnitines accumulation related to defective β-oxidation in the mutant cells. e) A heatmap showing the relative abundance of long chain fatty acids in control and *Pkd1* mutant MEFs. The panel shows that there is no clustering of LCFA levels by genotype, though some are generally lower in mutants than in controls. f, g) Boxplots showing levels of specific fatty acids: C16:0 (f) and C18:0 (g) in control and *Pkd1* mutant MEFs. h, i) Analysis of published lipidomic data obtained from *Pkd1*^*fll-*^; *Ksp-Cre* and control mouse kidneys [35]. Values were normalized to total lipid content in each sample to express as mol-percent data [35]. A plot showing phosphatidylcholine ether phospholipid (PC(O-)) levels in kidney tissues of *Pkd1* mutant mice compared to those of control mice (h). A plot showing levels of phospholipids containing VLCFA (C27:1) in kidney tissues of control and *Pkd1* mutant mice (i).

Taken together, these data suggest peroxisome function is preserved in ADPKD.

### Evaluating peroxisome lipid metabolism in published lipidomic and metabolomic studies of Pkd1 mutant cells and kidneys

To evaluate the generalizability of our findings, we analyzed the data from a recently published study of a different set of *Pkd1* mutant cells and cystic kidneys [35]. We first reviewed the metabolomic data obtained from mouse embryonic fibroblast cells (MEF) with/without *Pkd1* activity [35]. As measured analytes were not the same as ours, we evaluated other markers of peroxisome activity. Peroxisomes play central roles in the biosynthesis of plasmalogen, a special group of phospholipids, and docosahexaenoic acid (DHA), and downregulation in the biosynthesis of these lipid species is indicative of a peroxisome disorder [36, 37]. We therefore compared the levels of plasmalogen C16:0, plasmalogen C18:0, and DHA in control and *Pkd1* mutant MEFs. We found no significant difference in the levels of either plasmalogen or DHA, suggesting peroxisomal lipid biosynthesis is preserved in *Pkd1* ^-/-^ MEFs (Figure 6a–c). In striking contrast, however, the *Pkd1*^*-/-*^ MEFs exhibit accumulation of multiple acylcarnitines consistent with a mitochondrial fatty oxidation defect that we and others had previously described (Figure 6d) [4, 35]. The results for LCFA were mixed. While some fatty acids were generally lower in the mutant MEFs compared to controls (Figure 6e–g), especially C18:0 as was seen in the *Pkd1*^*fl/fl*^; *Ksp-Cre* animals (Figure 5), for many others there was no clear pattern based on genotype.

A recent study identified characteristic changes in the phospholipid composition in a broad spectrum of peroxisome diseases including Zellweger syndrome, ALD and single enzyme deficiencies [38]. Phosphatidylcholine ether phospholipids [PC(O-)] are lower and phospholipids containing VLCFA (>C22 fatty acids in one fatty acid side chain) are higher in fibroblasts of individuals with peroxisomal disorders. We therefore analyzed the published phospholipid profiles of *Pkd1*^*fll-*^; *Ksp-Cre* mice [35] and confirmed that total levels of PC(O-) and phospholipids containing VLCFA were not significantly different in cystic kidneys (Figure 6i). Lastly, in agreement with a previous report, we observed glucosylceramide (hexosylceramide) accumulation in the cystic kidneys [39], accompanied by downregulation of sphingomyelin synthesis (Supplementary Figure S3).

In conclusion, the results from re-analysis of published independent datasets of *Pkd1* mutant cells and kidneys are consistent with our findings and strongly argue against significant peroxisome dysfunction as a primary contributor to the pathogenesis of ADPKD and instead point to defective mitochondrial fatty acid oxidation as we had previously reported.

### Imaging studies to compare the localization of peroxisomal proteins and Polycystin-1

We have recently shown that a C-terminal cleavage product of PC1 (PC1-CTT) traffics to mitochondria and likely influences mitochondrial fusion/fission [6]. We therefore wondered if PC1-CTT would co-localize with PEX3 in portions of mitochondria that contribute to peroxisome formation [12], thereby ultimately localizing in peroxisomes.

Using live cell imaging studies in NIH3T3 cells with exogenous expression of PC1 CTT-mCherry and either PEX3 or peroxisome-targeted EGFP-SKL, we found PC1-CTT co-localized with mitochondrial markers and sometimes adjacent peroxisomes, but not to peroxisomes distant from mitochondria (Figure 7a and b). In fact, quantification of those images revealed three distinct patterns of PC1-CTT localization: a large group exclusively in mitochondria; a smaller proportion in areas with co-localized mitochondrial and peroxisomal markers; and a very small subset in mature peroxisome (EGFP-SKL positive vesicles) restricted to areas where mitochondria and peroxisome were adjacent, suggesting PC1-CTT is continuously localized in mitochondria and the PC1 product does not traffic to peroxisomes (Figure 7a and c). We also saw no specific co-localization of PC1 fragments with catalase in an mIMCD3 cell line with stable expression of full-length HA-tagged PC1 (Figure 7d). Taken together, we did not see evidence suggestive of peroxisomal targeting of the full-length PC1 or the PC1 cleavage product.

**Figure 7.**
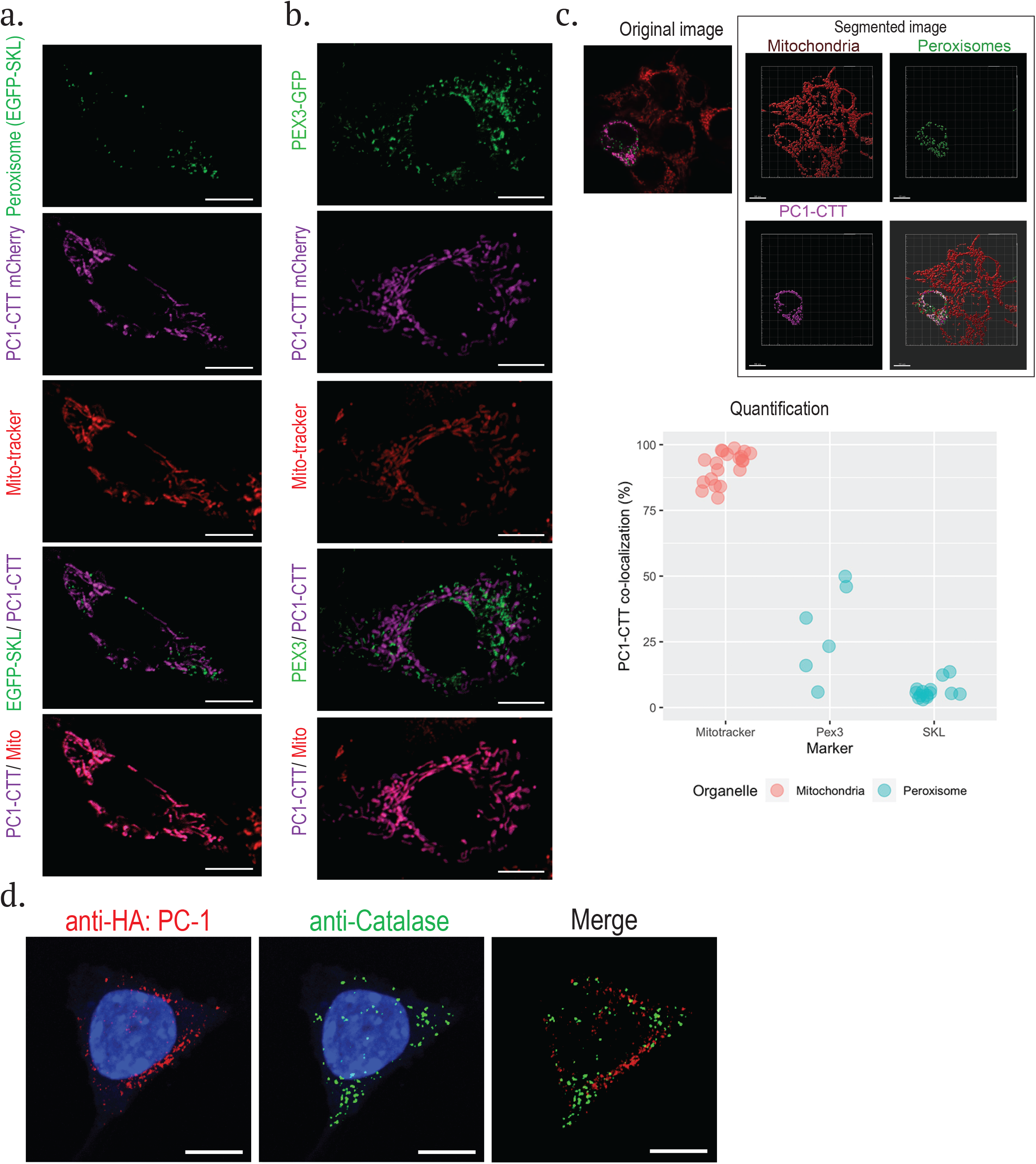
Co-localization study of PC1-CTT and peroxisomal proteins: EGFP-SKL or PEX3-GFP. a) Representative images of live cell imaging for a peroxisomal marker (EGFP-SKL, green), PC1-CTT (mCherry, magenta) and mitochondria (MitoTracker-Deep-Red, red) in NIH3T3 cells. Scale bar, 10µm. b) Representative images of live cell imaging for GFP-PEX3 (green), PC1-CTT (magenta) and MitoTracker (red) in NIH3T3 cells. Scale bar, 10µm. c) An example of image segmentation to quantify the PC1-CTT colocalization (original image: a top-left panel, segmented images: middle and right panels). The image quantification shows three distinct classification of PC1-CTT localization (a bottom panel) suggesting PC1-CTT does not traffic to peroxisomes. d) Representative images of double fluorescent staining for an HA-tagged full-length PC1 (red), the peroxisomal marker catalase (green) in mIMCD3 cells. FL-PC1-HA showed no obvious peroxisomal pattern. Scale bar, 10µm.

## Discussion

Multiple studies suggest that metabolic reprogramming is an intrinsic property of cells lacking *Pkd1*, although its precise contribution to cystogenesis remains unclear [2-4, 35]. Recent studies have further shown that targeting metabolism can be an effective strategy for treating orthologous/non-orthologous ADPKD models by modulating mitochondrial activity with pharmacological interventions or ketosis-inducing therapies [40, 41]. Given the therapeutic potential of this approach, we have sought to determine the mechanisms responsible for producing the altered cellular metabolic state. A number of recent studies have focused on the mitochondrion, with multiple groups reporting structural and functional abnormalities in cells and tissues of *Pkd1* mutants [4-7, 42]. We and others have also described direct physical and functional links between PC1 and mitochondria [6, 7, 42].

Despite the strong evidence linking PC1 to mitochondria, cystic disease is not a feature of mitochondrial diseases. In contrast, renal cystic disease is a feature of some inherited peroxisomal disorders, though the underlying mechanisms remain undefined [13]. Peroxisomes function as close collaborators of mitochondria in fatty acid metabolism and in regulation of the response to oxidative stress [10]. Furthermore, patients with Zellweger syndrome and *Pex5* mutant mice, a disease model for Zellweger syndrome, have altered mitochondrial morphology implicating a close pathophysiological interaction between the two organelles [15, 17].

In this study, we hypothesized that a peroxisomal defect might be an upstream factor that causes the mitochondrial defects observed in ADPKD. We examined peroxisome properties including its biogenesis pathway under normal and stressed conditions (Figures 1) and its function in fatty acid metabolism in ADPKD models (Figures 4 and 5) but found no consistent pattern of abnormalities. We also examined whether the C-terminal cleavage product of PC1 (PC1-CTT), which we had previously shown localizes to mitochondria, might also traffic to peroxisomes as had been reported for PEX3, a mitochondria-peroxisome dual targeting protein [12] (Figure 7). Regardless of which marker we used (an endogenous peroxisomal protein and PEX3, EGFP tagged with a peroxisomal targeting sequence, or EGFP-tagged PEX3), we found minimal co-localization with either PC1-CTT or full-length PC1.

We also queried published metabolomic and lipidomic datasets for evidence of peroxisome-related abnormalities (Figure 6) [35]. While we did not find robust evidence in support of this hypothesis, the data provide further evidence of mitochondrial fatty acid oxidation defects. Acylcarnitine levels were increased, and increased acylcarnitine levels are a readout for certain classes of defective mitochondrial fatty acid oxidation [43, 44]. This finding is also consistent with our earlier report of increased urinary acylcarnitines in the urine of *Pkd1* mutant mice [1]. Increased urinary excretion of acylcarnitines may be a mechanism by which excess acyl groups, unable to be adequately metabolized, are removed from the body [44].

One unexpected finding was that LCFA levels were reduced in the kidneys of *Pkd1* mutants (Figure 5) and *Pkd1* deficient MEFs (Figure 6) compared to controls. While the difference in the cystic kidneys is likely in part due to dilutional effects of cyst fluid contributing to total kidney weight, that cannot explain the differences observed in the mutant MEFs. The significance of this finding is unclear, but it further suggests that dysregulated lipid metabolism is a consequence of *Pkd1* mutation.

In sum, our studies have yielded consistent results showing that loss of *Pkd1* does not disrupt peroxisome biogenesis nor peroxisome-dependent fatty acid metabolism. These data refute our hypothesis that the previously reported mitochondrial abnormalities might in part be a downstream consequence of primary peroxisomal dysfunction and instead further suggest that mitochondrial dysfunction is a direct effect of PC1 loss. Finally, peroxisomes are organelles which possess indispensable functions not only in lipid metabolism but also in ROS regulation [45], and the innate immune system [46], two processes previously implicated in ADPKD pathogenesis [47, 48]. The current study cannot exclude the possibility that these other peroxisomal functions might be dysregulated in ADPKD and thus contribute to its pathogenesis, but our results suggest that any connection between PC1 and peroxisome function is likely to be indirect.

## Ethics Statement

All studies were performed using protocols approved by NIH Animal Care and Use Committee as appropriate, and mice were kept and cared in pathogen-free animal facilities accredited by the American Association for the Accreditation of Laboratory Animal Care and meet federal (NIH) guidelines for the humane and appropriate care of laboratory animal.

## Materials and Methods

### Cell lines

Kidney cell lines derived from *Pkd1* ^*fl/f*^ animals (respectively named as 121112C LTL, 94414 LTL and 96784 LTL cells) were established as previously described [4]. Cells were cultured in a humidified atmosphere of 5% CO2 at 37°C with Dulbecco’s minimal essential medium and Ham’s F12 nutrient mixture media (DMEM/F12) (Fisher Scientific, 15-090-CV) containing 2% fetal bovine serum (FBS) (GEMINI Bio, 100-106), 1?×?Insulin-Transferrin-Selenium (Thermo Fisher Scientific, 41400-045), 5?µM dexamethasone (SIGMA, D1756), 10?ng/ml EGF (SIGMA, SRP3196), 1?nM 3,3′,5-Triiodo-L-thyronine (Sigma-Aldrich, T6397), 1?×?GlutaMax (Thermo Fisher Scientific, 35050-061) and 10?mM HEPES (Quality Biological, 118-089-721). Since the parental cell line, 94414 LTL, is derived from a *Pkd1*^*flox/flox*^ mouse also expressing the Luo reporter, it expresses a GFP or RFP fluorescent protein based on the genotypes, only two pairs of cell lines were used for the imaging studies (121112 LTL. 96784 LTL). The NIH3T3 cell line was maintained with DMEM media (Fisher Scientific, 10-017-CV) containing 10% FBS. A Flp-In mIMCD3 cell line stably expressing C-terminal HA-tagged human PC1 and control cells were established as previously described [6] following the manufacturer’s instructions and maintained with DMEM/F12 media containing 10% FBS.

### Immunofluorescence

Cultured cells were seeded on glass coverslips (Neuvitro, H-22-1.5-pdl or GG-22-1.5-collagen). After fixation with 4% formamide PBS, the cells were permeabilized with 0.1 % or 0.5 % Triton X-100 (Sigma-Aldrich, T9284) in PBS for 10 minutes at room temperature. The cells were incubated with an Odyssey Blocking Buffer (LI-COR, 927-40000) for 30 minutes, then incubated with a rabbit polyclonal anti-PMP70 (Thermo Fisher Scientific, PA1-650, 1: 200), a goat anti-catalase (R&D systems, AF3398, 1: 150), a rabbit anti-Tom20 (Santa Cruz Biotechnology, FL-145, 1: 100), or a mouse anti-HA (MBL, M180-3, 1: 200) antibody followed by incubation with fluorescently labeled secondary antibodies (Thermo Fisher Scientific, [A-31572, 1:1000], [A-27034, 1:1000], [A-28180, 1:1000], [A-32814, 1:1000]). The stained coverslips were mounted onto microscope slides (Fisher Scientific, 12-550-15) with a DAPI-containing mounting media (Electron Microscopy Sciences, 17984-24). Image data were acquired using a confocal microscope (LSM780, Zeiss) and were visualized using ImageJ [49], and segmented and quantified using Imaris (Bitplane).

### Peroxisome temperature sensitivity test

Control and *Pkd1*^*-/-*^ mouse kidney epithelial cells were seeded on glass coverslips (Neuvitro, H-22-1.5-pdl or GG-22-1.5-collagen) and the cells were cultured for at least 16 hours at 37 °C. To test for a peroxisome defect under high temperature conditions, the cells were moved into a humidified atmosphere of 5% CO2 at 40 °C and the cells were cultured for 48 hours. After incubation at either temperature, the cells were washed twice with PBS, then fixed with 4% formamide PBS, followed by permeabilization with 0.5 % Triton X-100 (Sigma-Aldrich, T9284) in PBS for 10 minutes at room temperature followed by immunostaining for PMP70/catalase and image acquisition. The experiments were done twice, and at least 10 fields were randomly taken in each condition for the quantification.

### Immunoblotting

Culture cells were lysed with a RIPA lysis buffer (EMD Millipore, 20-188) containing a cOmplete Protease Inhibitor Cocktail (Sigma-Aldrich, 4693116001). Protein concentration was determined using a Pierce BCA Protein Assay Kit (Thermo Fisher Scientific, 23225), then the samples were reduced by a Sample Reducing Agent (Fisher Scientific, NP0009). Prepared samples were applied to 4-12% gradient Bris-Tris gels (Thermo Fisher Scientific, NP0335) and electrophoresed. Protein transfer was performed by an iBlot blotting system (Thermo Fisher Scientific) and nitrocellulose membranes (Thermo Fisher Scientific, IB301002). The membranes were immunologically stained with rabbit anti-PEX3 (Proteintech, 10946-1-AP, 1: 500), rabbit anti-PEX5 (Proteintech, 12545-1-AP, 1: 500), rabbit anti-PMP70 (Thermo Fisher Scientific, PA1-650, 1: 500) or mouse anti-α-Tubulin (Abcam, ab7291, 1: 20,000) antibodies followed by labeling with secondary antibodies. The specific band for each protein was detected and quantified by an Odyssey Infrared Imaging System (LI-COR). For quantification, signal intensity of each peroxisome-associated protein was measured using the Odyssey Infrared Imaging System and the obtained value was normalized to the intensity of α-tubulin band detected in the same lane. The experiments were done three independent times.

Plasmids

For the PC1-CTT-mCherry expression plasmid, the PC1-CTT (mPC-4119) [6] insert was fused with mCherry and cloned into pcDNA3 using an In-Fusion HD Plus kit (Takara Bio, 638909). The expression plasmids for EGFP-SLK (pEGFP-C1+SKL) and PEX3-GFP (pERB264) were produced by Jay Brenman (Addgene plasmid # 53450; http://n2t.net/addgene:53450; RRID:Addgene_53450) [28] and Michael Lampson (Addgene plasmid # 67764; http://n2t.net/addgene:67764; RRID:Addgene_67764) [50] respectively.

### Live cell imaging

Cells were seeded on glass coverslips (Neuvitro, H-22-1.5-pdl) or culture dishes with a polymer coverslip bottom (ibidi, 81156), then the cells were transfected with plasmids using a LipoD293 DNA transfection reagent (SignaGen Laboratories, SL100668) at 24–48 hours before the imaging studies. For investigating peroxisome matrix import, EGFP-SKL was transfected into *Pkd1* control and mutant kidney cells. The experiments were done at least twice, and a total of approximately 1,000 to 1,400 SKL-EGFP positive vesicles/cell line-genotype were processed by ImageJ [49] or Imaris (Bitplane). Ellipticity was measured by calculating the minor axis/(minor axis + major axis) of each vesicle using Imaris. For PC1-CTT localization studies, NIH3T3 cells were transfected with a plasmid expressing PC1-CTT-mCherry together with a plasmid expressing either EGFP-SKL or PEX3-GFP. Nucleus and Mitochondria were visualized by Hoechst 33342 (Thermo Fisher Scientific, H3570) and MitoTracker Deep Red FM (Thermo Fisher Scientific, M22426) respectively. Image data were obtained with a confocal microscope (LSM780, Zeiss), and the obtained images were visualized using ImageJ [49]. The images were segmented and quantified by Imaris using the Surface tool. The experiments were done at least three times, and at least 10 fields were randomly taken in each condition for the quantification.

### Long chain fatty acids assay by liquid chromatography–mass spectrometry (LC/MS)

High-performance liquid chromatography (HPLC) grade solvents were purchased from Sigma-Aldrich.

### Peroxisome β-oxidation study

Docosanoic-22, 22, 22-d3 acid (CDN isotope, D-5708) was dissolved in dimethyl sulfoxide (DMSO) for preparation of 6mM stock solution. Mice kidney epithelial cells cultured in 100mm culture dishes were incubated with 60µM of docosanoic-22, 22, 22-d3 acid (D3-C22:0) dissolved in culture media. After 48 hours, the cells were washed with phosphate buffered saline (PBS) twice and detached by a Trypsin:EDTA solution (GEMINIBio, 400-151), then the cells were collected into centrifuge tubes followed by centrifugation at 1000g for 5 minutes. Cell pellets were further washed with PBS followed by centrifugation at 1000g for 5minutes. The collected cell pellets were saponified by adding 300µL of 0.4M NaOH in MeOH and vortexed at room temperature overnight, then mixed with 20µL formic acid and centrifuged. The supernatant (250µL) was transferred into an LC-MS vial and diluted with 1mL MeOH. Total long chain fatty acids were analyzed using a Waters Acquity i-class UPLC coupled with a QExactive high resolution accurate-mass spectrometer (HRAM-MS, Thermo Scientific, Waltham, MA) with heated electrospray ionization (HESI-II) in negative ion mode. A Waters Cortecs UPLC T3 column (2.1mm x 100mm, 1.6µm) was used. The column was maintained at 40°C and the samples were kept in the autosampler at 4°C. The injection volume was 1µL. Chromatographic conditions were as follows: Solvent A: H_2_O, 0.01% AcOH; Solvent B: 62% ACN, 33% IPA, 5% H_2_O. The flow rate was 500µL/min, the run was isocratic with 93% B and the total running time was 7 min. Samples were analyzed in duplicate. Detection was based on *m/z* at the (M-H)^-^ for each LCFA. The ratios of D3 and natural fatty acids were calculated. The raw datasets are available in Supplementary Table 1.

### Mouse samples

*KSP-Cre; Pkd1*^*fl/fl*^ mice were bred as previously described [4]. Briefly, C57/BL6 *Pkd1*^*tm2Ggg*^ mice were crossed to the reporter mice C57/BL6 congenic B6.129S4-*Gt(ROSA)26Sor*^*tm1Sor*^*/J* (Jackson Laboratories, stock 003474) and to *Ksp-Cre B6*.*Cg-Tg(Cdh16-cre)91Igr/J* (Jackson Laboratories, stock 012237). A total of 35 kidney tissues (*Ksp-Cre* negative: n = 13, *Pkd1*^*fl/+*^*;Ksp-Cre*: n = 10 and *Pkd1*^*fl/fl*^*;Ksp-Cre*: n = 12) were harvested from mice at age of P3 or P6 and both total body weight and kidney weights were determined for each mouse. For the quantification of fatty acid levels in the tissues, each kidney sample was weighed. The 11 standard stocks were prepared by serial dilution of GLC 401 (Nu-Chek Prep, MN) in methanol and were kept at 4°C. The internal standard solution was prepared in methanol containing 1 µg/mL of [U-^13^C]-palmitic acid. Calibration standards were prepared by adding 100µL standard stocks, 100µL H_2_O and 400µL internal standard solution. The kidney samples (10-45mg) were prepared by hand-held micro-homogenizing tissue with 100µL MeOH, 100µL H_2_O and 400µL internal standard solution. Both calibration standards and kidney samples were saponified by adding 400µL of 0.4M NaOH in MeOH and kept at 55°C for 2h, then centrifuged and an aliquot of the supernatant (100µL) was transferred into an LC-MS vial, diluted with 0.9mL MeOH and mixed with 10µL formic acid. Total long chain fatty acids were detected and quantified utilizing a Thermo Scientific Vanquish UPLC with a Thermo Scientific Altis triple quadrupole mass spectrometer, heated electrospray ionization (HESI-II) in negative ion mode. A Waters Cortecs UPLC T3 column (2.1mm x 100mm, 1.6µm) was used. The column was maintained at 40°C and the samples were kept in the autosampler at 4°C. The injection volume was 1µL. Chromatographic conditions were as follows: Solvent A: H_2_O, 0.01% AcOH, Solvent B: 62% ACN, 33% IPA, 5% H_2_O. The flow rate was 500µL/min, the gradient was initially at 70% B, increasing to 95% B at 4.5 min, maintained 95% B to 7.1 min, then returned to 70% B at 7.4 min, the total running time was 8 min. Calibration standards and mouse kidney samples were analyzed in duplicate. Detection and quantification were based on *m/z* at the (M-H)^-^ with the result weight corrected for sample mass. The raw datasets are available ins Supplementary Table 2.

### Statistical analysis and data visualization

Statistic tests and data visualization were performed by software R (Version 3.6.1) [https://www.R-project.org/] running in the RStudio [http://www.rstudio.com/] and using the packages tidyverse [https://CRAN.R-project.org/package=tidyverse], psych [https://CRAN.R-project.org/package=psych], and pheatmap [https://CRAN.R-project.org/package=pheatmap]. All presented values are described as mean ± standard deviation. For the immunoblotting analysis, the quantified values were analyzed using paired t-test for comparison of matched pairs of cell lines. For the analysis of peroxisome β-oxidation study, results for three to five technical replicates were averaged for each cell line and genotype. The corresponding three pairs of means were analyzed using paired t-test. Comparison of fatty acid concentrations between control kidneys and kidneys of *Pkd1* mutant mice was done using Student’s t test. For the analysis of published metabolomics and lipidomics, the data tables were obtained from published literature [35], and the datasets were processed using software R, and comparisons of the metrics between control and *Pkd1* mutant MEFs/mice were performed by Student’s t test.

## Acknowledgements

This work was supported in part by the following NIH intramural cores (NIDDK Advanced Light Microscopy and Image Analysis and NIDDK Clinical Mass Spectrometry). We thank Jeff Reece for his expert advice for imaging studies. Some parts shown diagrams in Fig. 1 and 4 were made using a template modified from a TogoPictureGallery (http://togotv.dbcls.jp/togopic.2016.3.html), freely distributed under a 2016 DBCLS TogoTV.

## Authors Contributions

Dr. Menezes generated cell lines; Dr. Terabayashi prepared cells for the temperature studies and the other imaging studies and performed microscopy experiments. Dr. Terabayashi prepared samples and performed immunoblotting. Dr. Zhou managed the animal colony and collected data. Dr. Terabayashi prepared the samples for the peroxisome β-oxidation assays, and Drs. Cai, Garraffo, and Walter performed mass spectrometry procedures and analyzed the obtained data. Drs. Terabayashi and Menezes analyzed R, ImageJ, and Imaris-based analyses. Dr. Terabayashi prepared figures; Dr. Terabayashi designed the studies, and Drs. Menezes and Germino supervised the study, and interpreted the data. Drs. Terabayashi, Menezes, and Germino wrote the manuscript. Drs. Cai, Garraffo and Walter wrote the methods for LC/MS in the Materials and Methods section.

