## Supplementary Figures for "*Pkd1* mutation has no apparent effects on peroxisome structure or lipid metabolism"

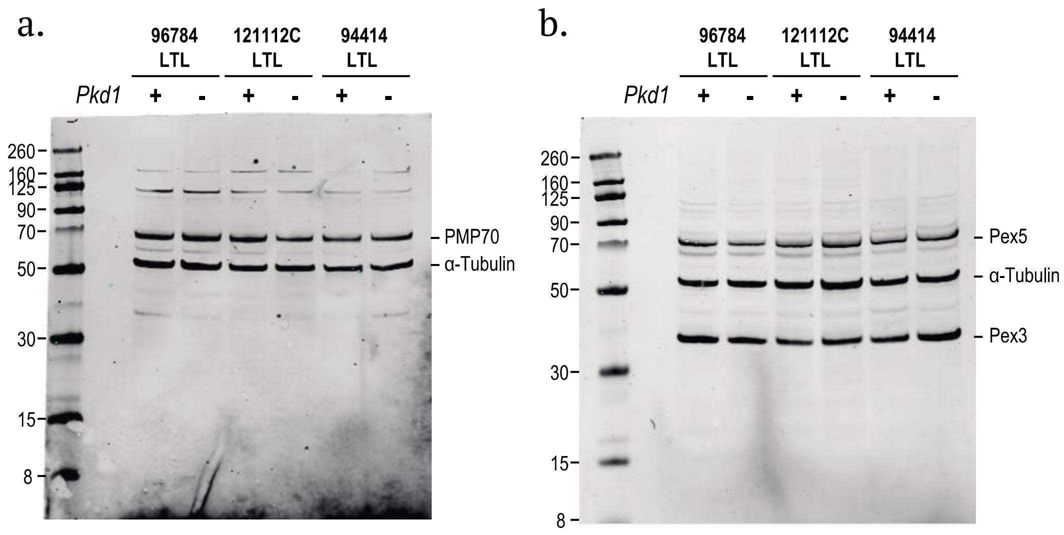


Figure S1.

**<u>Figure S1. Original immunoblots for Figure 2.</u>**

a, b) Original immunoblots for peroxisomal associated proteins: PMP70 (a), Pex3 and Pex5 (b) in 3 different pair of kidney cell lines (wild type and *Pkd1* mutant cells of 96784LTL, 121112C LTL, and 94414LTL cells). α-tubulin was used as a loading control.

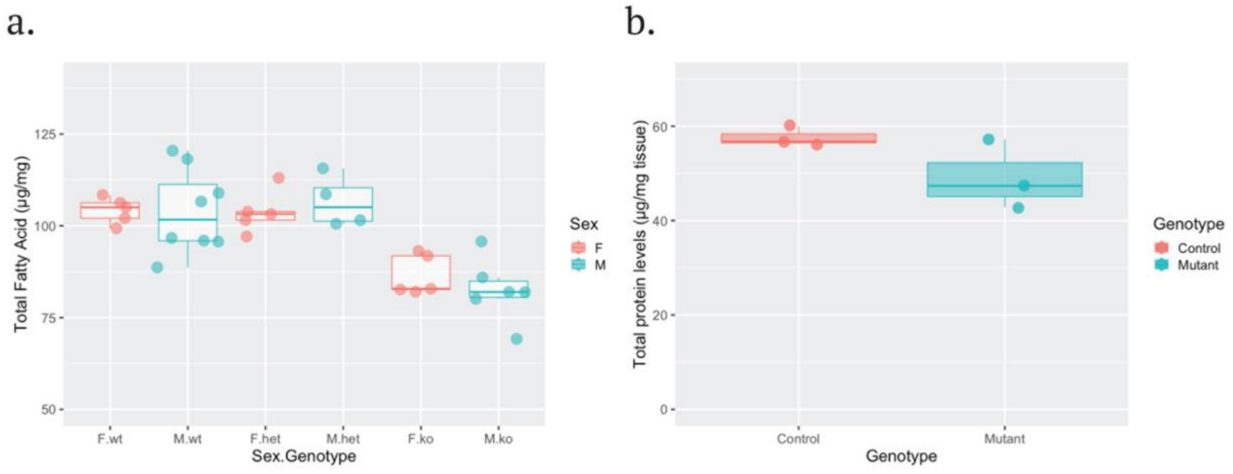


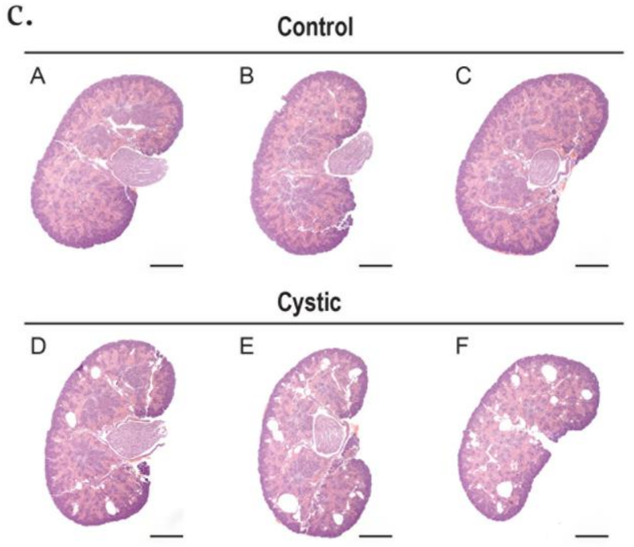


| Sample | Genotype | BW (g) | LKW (g) | RKW (g) | KBW | Protein Conc. (µg/mg tissue) |
|---|---|---|---|---|---|---|
| A | Control | 1.72 | 0.0101 | 0.0101 | 0.0059 | 60.0 |
| B | Control | 1.88 | 0.0095 | 0.0109 | 0.0054 | 56.7 |
| C | Control | 1.98 | 0.0116 | 0.0126 | 0.0061 | 56.2 |
| D | Mutant | 1.71 | 0.0113 | 0.0126 | 0.0070 | 57.2 |
| E | Mutant | 1.68 | 0.0122 | 0.0117 | 0.0071 | 42.8 |
| F | Mutant | 1.81 | 0.0124 | 0.0132 | 0.0071 | 47.4 |

Figure S2.

**Figure S2. Sex does not affect the total fatty acids level in ADPKD model mice.**

a) A boxplot presenting comparison of sex/genotype (x-axis) and the level of total fatty acids (C10–C24) (y-axis) of control (wt: *Ksp-Cre* negative (female n = 5, male n = 8), het: *Pkd1*$^{fl/+}$*;Ksp-Cre* (female n = 5, male n = 4)) and *Pkd1*$^{fl/fl}$*;Ksp-Cre* mice (female n = 5, male n = 6) implicated no difference between male and female within a genotype (ANOVA, p = 0.78). Mice with sex unknown (het: n = 1, ko: n=1) were excluded from the analysis.

b) A boxplot shows total protein levels (x-axis) of control (*Ksp-Cre* negative, n = 3) and mutant (*Pkd1*$^{fl/fl}$*;Ksp-Cre*, n = 3) mice (control 57.65 ±2.05 µg/mg vs mutant 49.13±7.35 µg/mg, p=0.13).

c) Overview of the control and the mutant mouse kidneys at postnatal day 3 on HE-stained sagittal sections (Scale bars, 500 µm.) presenting early cystic phase of *Pkd1*$^{fl/fl}$*;Ksp-Cre* mice (left panels). A table on the right panel presenting sample identifications (the alphabets corresponding to the ones labelled on the images of the kidneys on the left panels), body weight (BW), left kidney weight (LKW), right kidney weight (RKW), the kidney to body weight ratios (KBW), and tissue protein concentration.

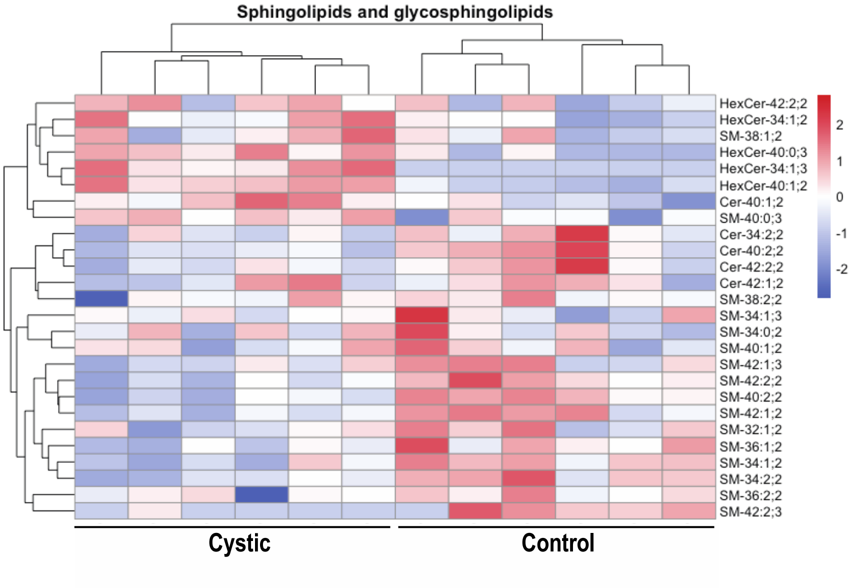


Figure S3.

**<u>Figure S3. Increased hexosylceramide in ADPKD mice model.</u>**

A heatmap showing sphingolipid and glycosphingolipid comparing kidneys of control and *Pkd1*$^{fl/-}$*; Ksp-Cre* mice. Consistent with a previous report (Natoli et al. 2010), clustering shows clear separation based on *Pkd1* genotype suggesting glucosylceramide (hexosylceramide: HexCer) accumulation together with downregulation of sphingomyelin (SM) synthesis in the cystic kidneys.
